# IL-12 signaling promotes TET2-mediated DNA demethylation during CD8 T cell effector differentiation

**DOI:** 10.1101/2020.11.02.365148

**Authors:** Caitlin C. Zebley, Hossam A. Abdelsamed, Hazem E. Ghoneim, Shanta Alli, Dalia Haydar, Tarsha Harris, Maureen A. McGargill, Giedre Krenciute, Ben Youngblood

## Abstract

CD8 T cell memory differentiation endows T cells with an ability to rapidly induce effector functions upon pathogen re-encounter. While it is well established that substantial epigenetic remodeling occurs during the effector stage of the immune response, the signaling events that imprint CD8 T cells with these stable epigenetic programs are not well-defined. To gain insight into the signaling determinants of effector-associated epigenetic programming among CD8 T cells, we explored the role of IL-12 in the imprinting of IFNg expression during human CD8 T cell priming. We observed that TCR-mediated stimulation of human naïve CD8 T cells is not sufficient to induce substantial demethylation of the IFNg promotor. However, TCR stimulation in the presence of the inflammatory cytokine, IL-12, resulted in significant and stable demethylation of the IFNg locus that was commensurate with an increase in IFNg expression. We further show that IL-12-associated demethylation of the IFNg locus is coupled to cell division through TET2-dependent passive demethylation in an *ex vivo* human CAR T cell model system and an *in vivo* immunologically competent murine system. Collectively, these data illustrate that IL-12 signaling promotes TET2-mediated effector epigenetic programming in CD8 T cells during the primary immune response and serve as proof of concept that signal 3 cytokines can be used to guide the induction of epigenetically regulated traits among T cells used for adoptive immunotherapies.

## INTRODUCTION

Adoptive transfer of T cells into cancer patients has proven efficacious against multiple tumor targets (1-3). Despite the irrefutable potential of this approach, many patients still succumb to their disease, prompting investigation into correlates of clinical response as well as methods to improve upon T cell-based therapies. IL-12 has been shown as an essential cytokine for enhancing the anti-tumor ability of T cells in multiple settings including intra-tumoral administration (4,5), facilitating successful anti-PD1 immunotherapy (6), and improving adoptive cell therapy approaches such as chimeric antigen receptor (CAR) T cell-mediated tumor eradication (7-9). IL-12-mediated tumoricidal activity has been directly linked to an upregulation in IFNg expression (10). Prior work in murine model systems has established that IL-12 signaling during T cell priming promotes IFNg expression in both effector and memory T cells and is critical for establishing long-lived functional memory CD8 T cells (11-13). Given its vital role in mediating a successful immune response, a deeper understanding of the molecular mechanisms underlying IL-12-mediated epigenetic programming of T cells is needed.

Immunological imprinting of effector programs in CD8 T cells occurs following naïve T cell antigen recognition and is tailored to the cytokine signals received from the innate immune system. Antigen presenting cells (APCs), such as dendritic cells, sense pathogens through toll like receptors which triggers production of proinflammatory cytokines including IL-12 (14,15). While naïve CD8 T cells are able to proliferate in response to TCR (signal 1) and CD28 (signal 2), signal 3 provided by either IL-12 or type I IFN is necessary for the development of a functional effector response (12). Importantly, IL-12 or type I IFN signals are not only crucial for the differentiation of naïve CD8 T cells into effector cells but are required for the development of memory cells (12). The critical role of IL-12 and type I IFN in memory formation is consistent with previous work in both humans and mice demonstrating that memory CD8 T cells originate from a subset of fate-permissive effector CD8 T cells (16). These memory CD8 T cells retain effector epigenetic programs which allow them to rapidly recall effector functions upon antigen reencounter (17). The ability of memory CD8 T cells to rapidly elicit effector functions contributes to their overall capacity to provide the host with life-long protection against previously encountered pathogens.

Persistence of memory CD8 T cells in the absence of their cognate antigen indicates that the cells undergo stable changes to gene regulation that can persist during homeostatic self-renewal (18). Dividing cells preserve transcriptionally repressive and permissive chromatin states through DNA methylation programs (19). Propagation of these acquired epigenetic programs preserves a population of long-lived CD8 T cells poised to recall an effector response. Indeed, our previous work shows that the effector DNA methylation programs of human memory CD8 T cells are conserved during antigen-independent self-renewal (20). DNA methylation programming is maintained by DNA methyltransferase 1 (DNMT1) which recognizes hemi-methylated CpG sites in order to re-establish 5-methylcytosine (5mC) on the daughter strand after DNA replication. DNA demethylation is driven by Ten-eleven translocation (TET) protein mediated oxidation of 5mC to 5-hydroxymethlcytosince and can occur passively through decreasing the affinity of DNMT1 binding resulting in the dilution of 5mC during DNA replication (21). While effector loci have broadly been shown to undergo demethylation during the priming stage of a T cell immune response, the mechanism governing acquisition of a memory T cell’s poised effector potential requires further investigation.

To better understand the molecular mechanisms reinforcing effector-associated epigenetic programs instilled during CD8 T cell differentiation, here we investigate the role of IL-12 in CD8 T cell priming with a focus on IFNg. Using a human chimeric anitgen receptor (CAR) model system, we report that IL-12 driven demethylation of the IFNg promotor is mediated by TET2 and occurs passively following cell division. We extend these results to an endogenous immune response using the well-established LCMV murine model of acute viral infection. Our results inform on the signaling events that promote effector-associated DNA demethylation during both human and mouse CD8 T cell effector differentiation and provide clinically relevant mechanistic insight regarding the optimization of adoptive T cell therapies.

## RESULTS

### IL-12 signaling promotes IFNg locus epigenetic reprogramming during human CD8 T cell effector differentiation

DNA methylation is known to play a crucial role in T cell differentiation. For instance, during effector differentiation the IFNg locus in CD8 T cells becomes demethylated commensurate with its high level of expression (22). To investigate the role of individual cytokines in effector differentiation-driven epigenetic reprogramming, we isolated naïve CD8 T cells from a healthy human donor and cultured them *in vitro* with anti-CD3/CD28 (TCR) plus different cytokines known to be crucial for CD8 T cell function including IL-2, IL-7, IL-12, IL-15, IL-18, and IL-21 **(Figure S1)**. While the majority of cytokines did not impact IFNg protein expression or locus methylation status, naïve CD8 T cells cultured in the presence of IL-12 and anti-CD3/CD28 exhibited both an increase in IFNg expression and underwent demethylation of the IFNg promotor prompting further investigation of IL-12 **(Figures 1A and S1)**. We next proceeded to define the kinetics for IFNg expression and locus demethylation. At the two, seven, and fourteen-day timepoints of the *in vitro* culture, we measured IFNg expression and examined the methylation status of the IFNg locus for naïve CD8 T cells stimulated with anti-CD3/CD28 plus IL-12. As early as day 2, the CD8 T cells cultured in the presence of IL-12 exhibited robust IFNg expression as compared to minimal IFNg expression seen in the CD8 T cells stimulated only with anti-CD3/CD28 **(Figures 1B - D)**. Further, the CD8 T cells stimulated with anti-CD3/CD28 and IL-12 maintained a heightened ability to express IFNg at both the seven and fourteen-day timepoints **(Figures 1B - D)**. Notably, the ability of these CD8 T cells to express IFNg was coupled to epigenetic changes at the IFNg locus. While unstimulated naïve CD8 T cells and naïve CD8 T cells stimulated with anti-CD3/CD28 remained methylated, naïve CD8 T cells stimulated with anti-CD3/CD28 in the presence of IL-12 underwent demethylation of the IFNg locus by day 7 and the demethylation persisted at day 14 **(Figure 1E)**. These data suggest that the presence of TCR signaling alone is not sufficient to fully upregulate IFNg expression or induce complete demethylation of the IFNg locus. The presence of proinflammatory cytokines, specifically IL-12, during CD8 T cell priming is necessary to drive the expression of IFNg and instill the effector-associated epigenetic programs of the IFNg locus.

**Figure 1.**
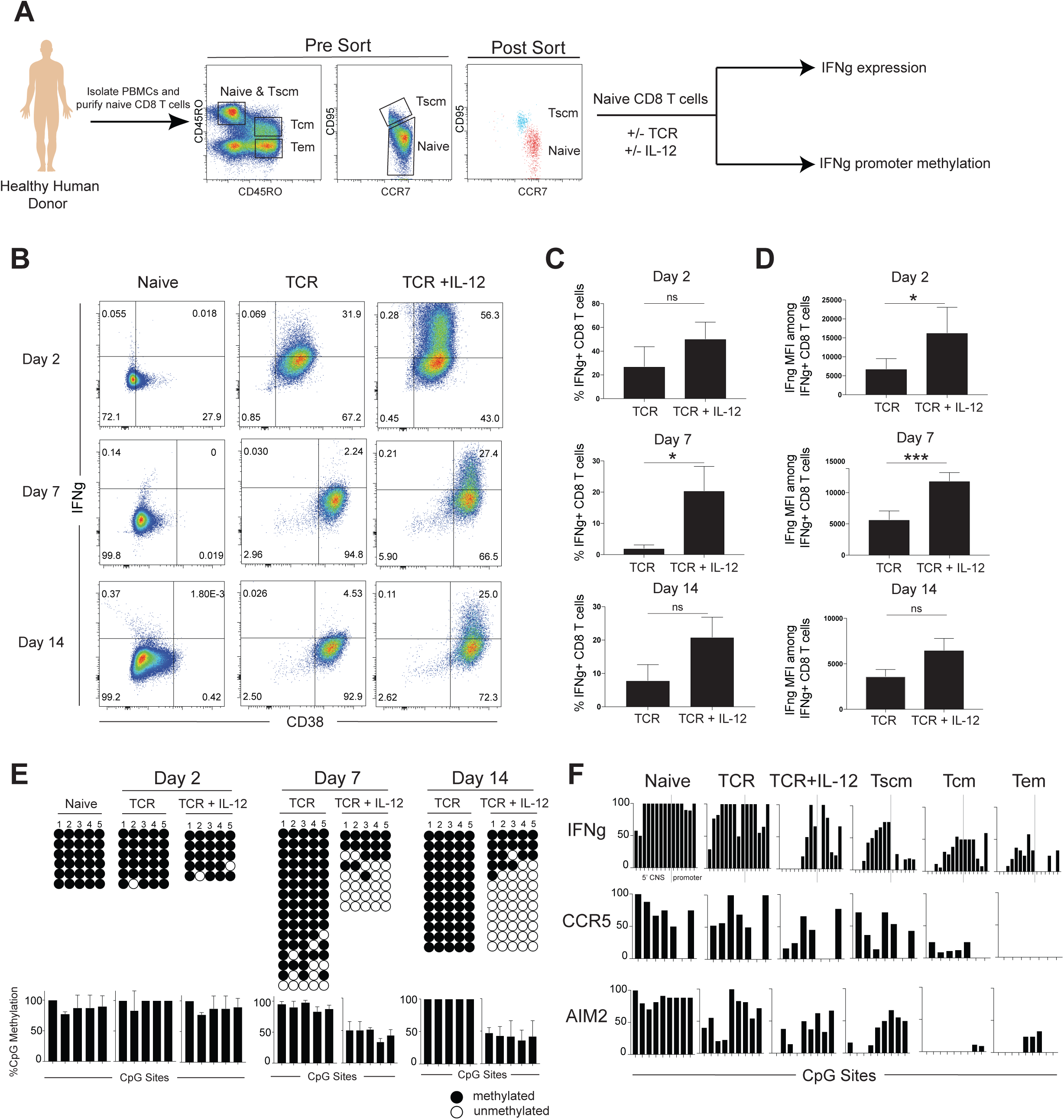
IL-12 signaling during human CD8 T cell priming promotes IFNg expression and locus demethylation. **(A)** Experimental setup showing isolation of naïve CD8 T cells for *in vitro* stimulation with or without anti CD3/CD28 (TCR) and with or without IL-12. **(B)** CD38 and IFNg expression of naïve, TCR stimulated, and TCR + IL-12 stimulated CD8 T cells after 2, 7, and 14 days. n=2 independent donors **(C)** Summary graph of the percentage of CD8 T cells expressing IFNg. **(D)** Summary graph of IFNg MFI among IFNg positive CD8 T cells. **(E)** Representative IFNg promotor bisulfite sequencing methylation profiles and summary graph of naïve, TCR stimulated, and TCR + IL-12 stimulated CD8 T cells after 2, 7, and 14 days. n=2 independent donors **(F)** Representative DMRs of genes known to be regulated by IL-12 derived from WGBS data of a healthy human donor.

Based on prior published work demonstrating that memory T cells transition through an effector stage of differentiation (16,23), we proceeded to ask whether IL-12 signaling induced DNA demethylation events were also present in freshly-isolated endogenous memory T cells from healthy human donors. Consistent with our loci-specific DNA methylation profiling, genome wide methylation profiling documented demethylation of the IFNg promotor as well as the 5’ conserved non-coding sequence (CNS) element (24) in the presence of TCR and IL-12 **(Figure 1F)**. Notably, these specific demethylation events were present in memory T cell subsets freshly isolated from healthy human donors (20,25). For further comparison, we looked at other genes (CCR5 and AIM2) (26,27) previously reported to be regulated by IL-12 and found similar demethylation events that were enriched in the long-lived endogenous T cell memory populations **(Figure 1F)**. Given that IL-12 enhances effector differentiation, our methylation data further support a model of memory T cell development whereby the memory CD8 T cells transition through an effector stage of the immune response.

### Cell division is coupled to epigenetic reprogramming of the IFNg locus during human effector CD8 T cell differentiation

Prior efforts to identify the origin of human memory CD8 T cells have demonstrated that long-lived memory T cells are derived from a subset of T cells that have undergone a proliferative burst during the effector stage of a primary immune response (23). Given that we did not observe notable demethylation of the IFNg locus until after 2 days of stimulation **(Figure 1E)**, we proceeded to evaluate whether cell division plays a role in establishing the observed epigenetic reprogramming. To evaluate the role of cell division in demethylation of the IFNg locus, CFSE-labeled naïve CD8 T cells were stimulated in the presence of anti-CD3/CD28 with and without IL-12. After one week, the CFSE-labeled CD8 T cells were assessed for IFNg expression. While both culture conditions resulted in significant cell division after stimulation with anti-CD3/CD28, the majority of IFNg expression was observed in those CD8 T cells that divided in the presence of IL-12 **(Figures 2A and 2B)**. We next FACS purified the CD8 T cells into undivided and divided cell populations to determine whether the IFNg expression was correlated with division-dependent changes in epigenetic programs. Bisulfite sequencing revealed that IFNg expression in the divided cell population was coupled to DNA demethylation of the IFNg locus. These results demonstrate that cell division promotes IFNg locus demethylation in CD8 T cells stimulated with anti-CD3/CD28 and IL-12 **(Figure 2C)**. While these results further support the conclusion that IL-12 signaling is critical for inducing IFNg expression, the association with cell division prompted us to further explore the enzymatic mechanism responsible for inducing DNA demethylation.

**Figure 2.**
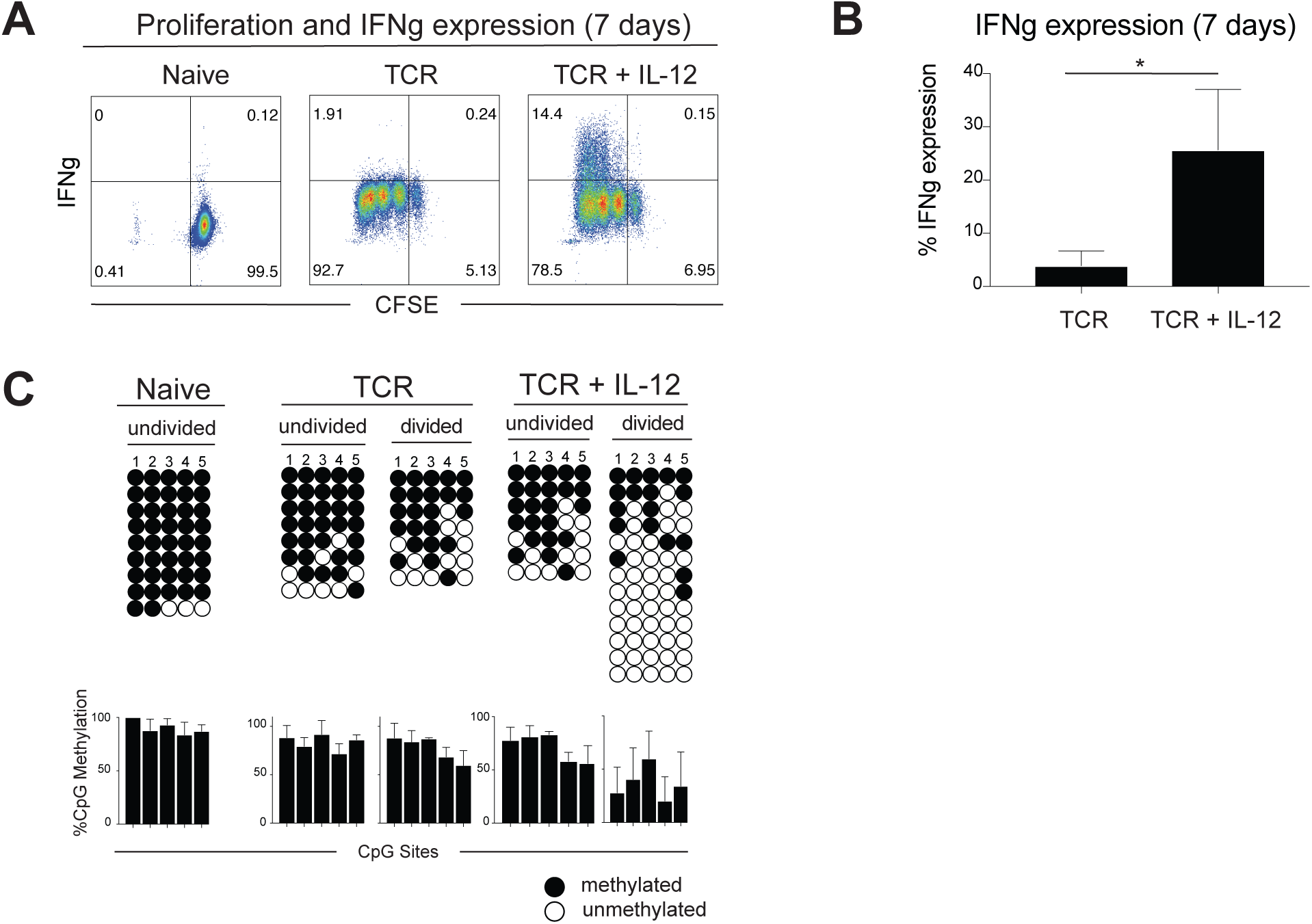
IL-12 mediated demethylation of the human IFNg promotor occurs after cell division. **(A)** Representative IFNg expression of CFSE labeled naïve, TCR stimulated and TCR + IL-12 stimulated CD8 T cells. n=4 independent donors **(B)** Summary of IFNg expression in CFSE labeled TCR stimulated and TCR + IL-12 stimulated CD8 T cells from panel A. n=4 independent donors **(C)** Representative IFNg promotor bisulfite sequencing methylation profiles and summary graph of naïve, TCR stimulated, and TCR + IL-12 stimulated CD8 T cells in FACS-purified undivided and divided cell populations after 7 days. Divided populations were sorted after 3 or more CFSE-defined divisions. n=2 independent donors

### IL-12 signaling promotes TET2 driven IFNg locus demethylation during human CAR T cell activation and expansion

Previous studies have shown that the ten eleven translocation (TET) enzymes oxidize 5-methylcytosine (5mC) to 5-hydroxymethylcytone (5hmC) (21). The maintenance methyltransferase, DNA methyltransferase 1 (DNMT1), does not readily recognize 5hmC which results in a failure to maintain DNA methylation after cell division, termed “passive” demethylation (21). Given our observation that IFNg demethylation occurs in a division-dependent manner, we used an *ex vivo* human CAR T cell system to ask whether TET2-mediated 5hmC deposition resulted in passive DNA demethylation of the IFNg locus. To investigate the role of TET2 in human CD8 T cell effector differentiation, human naïve CD8 T cells were FACS purified from a healthy donor, activated in the presence of anti-CD3/CD28 followed by electroporation with Cas9 and gRNA targeting either mCherry (control) or TET2 and subsequently transduced with a lentivirus that expressed the B7-H3-specific chimeric antigen receptor (CAR) (Haydar et al, unpublished). The cells were then expanded in the presence of IL-7 and IL-15 until day 10 at which point they were activated with plate-bound recombinant B7-H3 with or without IL-12 for 14 days prior to DNA methylation profiling **(Figure 3A)**. Loci specific bisulfite sequencing revealed that DNA demethylation of the IFNg locus was significantly enhanced by IL-12 as seen in the control KO, but strikingly IFNg locus demethylation did not occur in the absence of TET2 in conditions either with or without IL-12 **(Figure 3B)**. These results document the critical role of TET2 in driving demethylation of the human IFNg locus in the setting of inflammatory cytokines.

**Figure 3.**
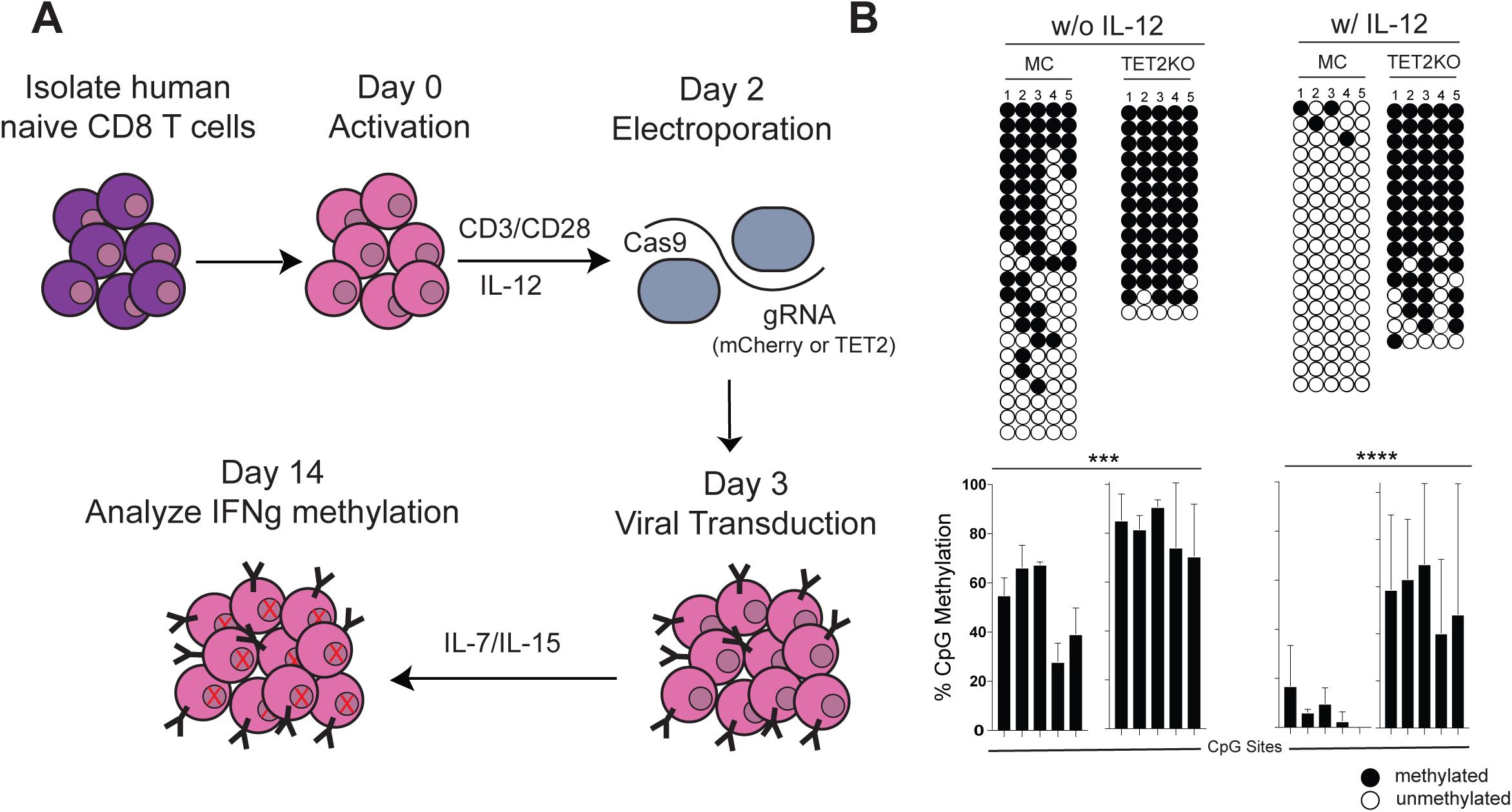
TET2 drives demethylation of the human IFNg promotor. **(A)** Experimental setup for TET2KO in activated human naive CD8 T cells. Human naïve CD8 T cells were FACs purified from a healthy donor, activated in the presence of TCR for 48 hours, electroporated with cas9 and gRNA targeting either mCherry (control) or TET2, transduced with lentivirus, expanded in the presence of IL-7 and IL-15 until day 10, and activated with plate-bound recombinant B7-H3 with or without IL-12 for 14 days prior to DNA methylation profiling. **(B)** Representative IFNg promotor bisulfite sequencing methylation profiles and summary graph of control or TET2KO *ex vivo* expanded CD8 T cells either in the absence or presence of IL-12. n=2 independent donors.

### *In vivo* bystander inflammation during T cell priming promotes TET2-mediated Ifng locus demethylation

We next sought to extend our findings by examining the role of inflammatory cytokines in an endogenous immune response using an immunocompetent model system. Previously we have demonstrated that demethylation of Ifng occurs in memory precursor T cells and persists into the development of all memory T cell subsets (16) **(Figures S2A & S2B)**. Building on these previous findings, we examined *Listeria monocytogenes* (LM)-specific effector CD8 T cells to determine the role of bystander inflammation on DNA methylation of the Ifng locus. LM-specific effector CD8 T cells undergo extensive proliferation during the effector response to acute LM infection. However, unlike acute lymphocytic choriomeningitis virus (LCMV)-specific effector CD8 T cells, they are resistant to demethylation of the Ifng 3’ CNS **(Figure S2C)**, a site which has been previously described as a regulatory element for Ifng expression (24). Notably, LM-specific effector CD8 T cells do undergo demethylation of the Ifng promotor (28) illustrating intergenic variability and focusing our efforts on identifying the signaling determinants for demethylation of the Ifng 3’ CNS.

Having identified a dichotomous methylation state among LM and LCMV-specific effector CD8 T cells, we sought to determine if the inflammatory milieu of an acute LCMV infection could promote further demethylation of the Ifng locus in LM-specific CD8 T cells. We proceeded to interrogate LM-specific CD8 T cells in the context of single infection with LM or co-infection with LM and acute LCMV. To track an LM-specific CD8 T cell response, we adoptively transferred ova-specific TCR transgenic OT-1 CD8 T cells into naive C57BL/6 mice prior to either infection with LM-OVA or co-infection with LM-OVA and acute LCMV **(Figure 4A)**. Relative to the OT-1 cells in mice infected only with LM-OVA, the OT-1 CD8 T cells exhibited a more activated phenotype in the context of LM-OVA and LCMV co-infection as exemplified by a higher percentage of KLRG1+CD127-cells **(Figure S2D)**. Having observed a difference in the phenotypes, we next FACS purified OT-1 effector CD8 T cells from mice either single infected with LM-OVA or co-infected with LM-OVA and LCMV. The OT-1 effector CD8 T cells remained mostly methylated at the Ifng 3’ CNS after acute LM-OVA infection. Remarkably, the OT-1 CD8 T cells become demethylated in the setting of co-infection with LM-OVA and acute LCMV **(Figure 4B)**. Demethylation of the Ifng 3’ CNS was maintained in OT-1 memory cells generated during coinfection with LM-OVA and acute LCMV **(Figure S2E)**, demonstrating the stability of DNA methylation programming instilled during effector differentiation. These differences in Ifng 3’ CNS methylation between LM-OVA and LM-OVA + LCMV generated OT-1 effector and memory CD8 T cells suggest a role for proinflammatory cytokines in providing the signals required to induce demethylation in the setting of co-infection.

**Figure 4.**
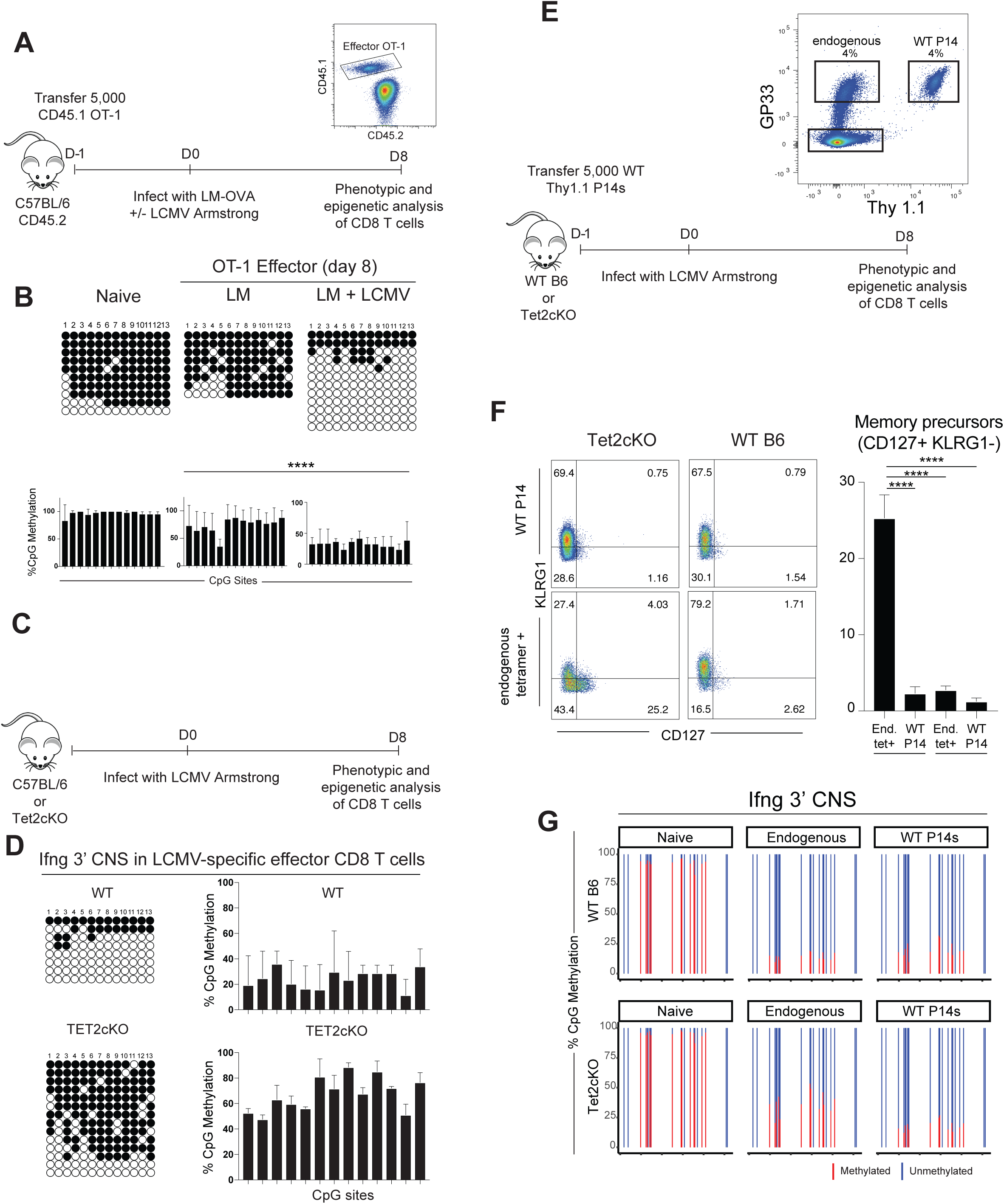
TET2 promotes inflammatory cytokine mediated demethylation of murine Ifng during effector CD8 T cell differentiation. **(A)** Experimental setup showing adoptive transfer of CD45.1 OT-1 CD8 T cells into WT B6 CD45.2 mice. One day later, the mice were infected with either LM-OVA or co-infected with LM-OVA and LCMV Armstrong. At the effector (8 day) and memory (>2 months) stages, CD45.1 OT1 CD8 T cells were FACS purified for phenotypic and DNA methylation analysis. **(B)** Representative and summary graphs of IFNg CNS bisulfite sequencing methylation profiles of naïve CD8 T cells and LM-specific and LCMV-specific memory CD8 T cells at day 28 post infection. n=6 for LM and n=4 for LM+LCMV summary graphs. **(C)** Representative bisulfite sequencing methylation profiles of the Ifng promotor and Ifng 3’ CNS in LCMV-specific effector CD8 T cells from WT or TET2cKO mice. **(D)** Experimental setup showing adoptive transfer of Thy1.1+ P14 CD8 T cells into Thy1.2+ WT B6 or TET2cKO mice. One day later, the mice were infected with LCMV Armstrong. At the effector (8 day) stages, endogenous and adoptively transferred GP33-specific CD8 T cells were phenotypically characterized and FACS purified for DNA methylation analysis. **(C)** Representative FACs analysis and summary graph characterizing the terminal effector (KLRG1+ CD127-) and memory precursor (KLRG1-CD127+) phenotype of either endogenous or adoptively transferred LCMV-specific CD8 T cells isolated from WT B6 or Tet2cKO mice. **(D)** Targeted methylation profiling of the IFNg 3’ CNS in naive, endogenous, and WT P14s CD8 T cells isolated from WT and Tet2cKO mice at the effector timepoint.

After establishing that the cytokine milieu resulting from LCMV infection enables demethylation of the Ifng 3’ CNS, we asked whether Tet2 was playing a role in this inflammatory-mediated process. To answer this question, we infected either WT or TET2cKO mice with acute LCMV and analyzed the LCMV-specific CD8 T cells at the effector timepoint. Bisulfite sequencing analysis revealed that the Ifng 3’ CNS was primarily demethylated in LCMV-specific CD8 T cells isolated from WT mice but remained mostly methylated in LCMV-specific CD8 T cells isolated from TET2cKO mice **(Figure 4C)**. To further investigate these differences in Ifng 3’ CNS methylation while controlling for environmental differences between mice, we transferred 5,000 WT Thy1.1 P14s into TET2cKO or B6 mice and infected the mice with LCMV Armstrong. From the same mouse, the transferred Thy1.1 P14s and endogenous LCMV-specific CD8 T cells were sorted at the effector timepoint **(Figure 4D)**. Phenotypic analysis revealed that the endogenous LCMV-specific CD8 T cells isolated from the TET2cKO mice had a significantly greater proportion of memory precursors (CD127+ KLRG1-) as compared to adoptively transferred P14 cells within the same animal **(Figure 4E)**. Conversely, endogenous LCMV-specific CD8 T cells from WT mice exhibited a similar phenotype to the adoptively transferred P14 cells within the same mouse **(Figure 4E)**. These results are consistent with a prior study examining the role of Tet2 in an LCMV T cell-mediated immune response (29). We next performed targeted DNA methylation profiling to interrogate the methylation status of the Ifng 3’ CNS. As expected, the endogenous naïve population retained methylation at the Ifng locus while the transferred Thy1.1 P14s underwent greater demethylation at the IFNg locus in both the WT and TET2cKO mice **(Figure 4F)**. Importantly, differences in methylation were observed among the endogenous WT and TET2cKO LCMV-specific effector CD8 T cells. While the endogenous LCMV-specific CD8 T cells underwent demethylation of the Ifng locus in the WT mice consistent with the demethylation observed in the transferred Thy1.1 P14s, this was not the case in the TET2cKO mice. The endogenous LCMV-specific CD8 T cells in the TET2cKO mice underwent only partial demethylation of the Ifng locus and remained significantly more methylated than the transferred Thy1.1 P14s isolated from the same mouse **(Figure 4E)**. This result demonstrates that TET2 regulates cytokine-mediated Ifng 3’ CNS demethylation during an *in vivo* CD8 T cell effector response to a viral infection. Collectively our results provide further mechanistic insight into how memory CD8 T cells acquire effector-associated epigenetic programs during the proliferative burst of an immune response.

## DISCUSSION

Here we show that IL-12 acts as a signal 3 cytokine during effector and memory CD8 T cell differentiation to drive TET2-mediated DNA demethylation of the IFNg locus during T cell priming. Collectively, our work data broadly demonstrates that communication between the innate and adaptive immune systems via cytokine signaling can lead to stable changes in DNA methylation. While T cells recognize antigenic peptide in the context of MHC and costimulation during activation, cytokine exposure introduces variability during the process of T cell differentiation. It has been previously shown that the presence of TCR signaling alone is not sufficient to produce a functional effector or memory CD8 T cell (13). The data we provide here highlight the importance of inflammatory cytokines in tailoring the epigenetic programs that are acquired during the priming stage of a human immune response that are maintained in long-lived memory T cells. Our results build upon previous work demonstrating that memory T cells arise from fate-permissive effector T cells and subsequently maintain effector-associated DNA methylation programs (16). Further, these observations are consistent with prior reports that memory T cells divide extensively after infection and are endowed with an effector epigenetic landscape which is maintained by quiescent memory cells (20,23).

Previous work has shown that T-bet requires signal transducer and activator of transcription 4 (Stat4) for complete IL-12-dependent development of murine T helper 1 (Th1) cells (30). Stat4 dimerization has been shown to regulate chromatin remodeling (31), yet Stat4 is only transiently activated and the mechanisms that dictate Th1 transcriptional regulation in effector and memory CD8 T cells have not been fully elucidated. While IL-12 is known to play a critical role in murine Th1 differentiation (32), here we show a direct link between signal 3 cytokine signaling and DNA methylation remodeling during human effector and memory CD8 T cell differentiation. Our data suggest that the extent of site-specific IFNg demethylation is regulated by TET2 with development of a memory precursor phenotype in TET2-deficient CD8 T cells. These results are consistent with previous studies reporting that loss of TET2 limited effector T cell differentiation (29). Our findings provide additional insight into recent reports showing improved clinical efficacy coupled to CAR T cells that had a mutation in TET2 and ultimately enriched for survival of TET2 KO CAR T cells with a central memory phenotype (33). Here we show that deletion of TET2 during naive to effector differentiation modifies the effector DNA methylation program in CAR T cells.

In order to incorporate mechanisms of memory T cell differentiation into the manufacturing process of CAR T cells, it is important to define when and how memory T cells acquire and maintain their effector potential. Consistent with prior investigations into the origin of human memory T cells, our work here shows that effector-associated epigenetic programs are acquired during the proliferative burst of an effector response and are promoted by signal 3 cytokines present during T cell priming. Future efforts to improve CAR T cell therapeutic approaches are now looking to exploit our understanding of memory T cell identification by incorporating cytokines into conditions for developing CAR T cells. Notably, with the field looking to expand current CAR T cell protocols in treating solid tumors, our work serves as a proof of principle that such engineering approaches may be implemented to enhance the effector potential of long-lived adoptive T cell therapies.

## MATERIALS AND METHODS

### Isolation and phenotypic analysis of mouse antigen-specific CD8 T cells

LM specific CD8 T cells were generated by adoptive transfer of ∼5000 naïve OT-1 TCR transgenic CD8 T cells (CD45.1/1+) into C57BL/6 mice (CD45.1/2+). Chimeric mice were subsequently infected with *Listeria monocytogenes* containing OVA by injection of 1.5×10^4 CFU per mouse. LCMV specific CD8 T cells were generated by adoptive transfer of ∼5000 congenically distinct naive P14 CD8 T cells. One day later, we infected the mice with acute lymphocytic choriomeningitis virus (LCMV). Acute LCMV infection was performed by i.p. injection of 2×10^5 PFU Armstrong strain per mouse. Chronic LCMV infections were performed by i.v. injection of 2×10^6 PFU LCMV per mouse using either Clone 13 strain. Antigen specific CD8 T cells were identified by gating on CD44hi CD8+ lymphocytes that were either tetramer+ or congenically labelled. Tetramer was obtained from the NIH Yerkes tetramer core facility. All antigen specific CD8 T cells were phenotypically analyzed by Flow Cytometry after surface staining using monoclonal antibodies for Thy1.1(BD clone OX-7), CD45.1 (Biolegend clone A20), CD45.2 (Biolegend clone 104), CD8 (Biolegend clone 53-6.7), Klrg1 (Biolegend clone 2F1), CD127 (Biolegend clone A7R34), CD44 (Biolegend clone IM7).

Tet2 floxed mice were obtained from Jackson laboratories (Catalog #017573) and crossed with granzyme B cre mice (34) to generate mice that conditionally delete Tet2 in activated CD8 T cells. Wild type C57B6 mice were purchased from Jackson laboratories.

### Isolation of human naïve CD8 T cells from healthy donor blood

This study was conducted with approval from the Institutional Review Board of St. Jude Children’s Research Hospital. Human peripheral blood mononuclear cells (PBMCs) were collected through the St. Jude Blood Bank, and samples for methylation profiling were collected under IRB protocol XPD15-086. PBMCs were purified from platelet apheresis blood unit by density gradient. Briefly, blood was diluted 1:2.5 using sterile Dulbecco’s phosphate-buffered saline (Life Technologies). The diluted blood was then overlayed above Ficol-Paque PLUS (GE Healthcare) at a final dilution of 1:2.5 (ficoll:diluted blood). The gradient was centrifuged at 400 xg with no brake for 20 minutes at room temperature. The PBMCs interphase layer was collected and washed with 2% fetal bovine serum (FBS)/1mM EDTA PBS buffer and then centrifuged at 400xg for 5 minutes.

### Flow cytometric analysis of human naive CD8 T cells

After enrichment of CD8 T cells, naive and memory CD8 T cell subsets were sorted using the following markers, as previously described (35,36). Naive CD8 T cells were defined as live CD8+, CCR7^+^, CD45RO^−^, CD45RA^+^, and CD95^−^ cells. CD8 Tem cells were defined as live CD8+, CCR7^−^, and CD45RO^+^ cells. T_CM_ cells were defined as live CD8+, CCR7^+^, and CD45RO^+^ cells. Tscm cells were defined as live CD8+, CCR7^+^, CD45RO^−^, and CD95^+^ cells. Naïve sorted cells were checked for purity (i.e., samples were considered pure if >90% of the cells had the desired phenotype). The naïve CD8 T cells were then cultured *ex vivo* with or without anti-CD3/CD28 (1:1 ratio) in the presence or absence of the following cytokines: IL-2 (10ng/mL), IL-7 (5ng/mL), IL-12 (10ng/mL), IL-15 (5ng/mL), IL-18 (10ng/mL), IL-21 (30ng/mL). At 2, 7, or 14 days, the levels of IFNg expression was examined by intracellular staining after exposure to 4 hours of GolgiStop and GolgiPlug (BD).

Human CD8 T cells were stained with the following antibodies: CCR7 (Biolegend clone G043H7), APCCy7 (Biolegend clone SK1), PeCy7 (Biolegend clone DX2), CD38 (Biolegend clone HB-7), IFNg (Biolegend clone 4S.B3) APC (Biolegend clone UCHL1).

### In vitro CD8 T cell ex vivo proliferation

Sorted naive CD8 T cells and memory CD8 T cell subsets were labeled with CFSE (Life Technologies) at a final concentration of 2 µM. CFSE-labeled cells were maintained in culture in RPMI containing 10% FBS, penicillin-streptomycin, and gentamycin. After 7 days of *ex vivo* culture at 37°C and 5% CO2, undivided and divided cells (third division and higher) were sorted and checked for purity (>90%). The levels of IFNg protein expression was determined by intracellular staining after exposure to 4 hours of GolgiStop and GolgiPlug (BD).

### Genomic Methylation Analysis

DNA was extracted from the sorted cells by using a DNA-extraction kit (Qiagen) and then bisulfite treated using an EZ DNA methylation kit (Zymo Research). The bisulfite-modified DNA-sequencing library was generated using the EpiGnome™ kit (Epicentre) per the manufacturer’s instructions. Bisulfite-modified DNA was PCR amplified using the following primers for mouse and human, respectively.

Mouse IFNg Forward: 5’-GTTTATTTTTATTGTTGTGGTTGGTAGCTG-3’

Mouse IFNg Reverse: 5’-CCTTTCTTCTCCAAATTACTTTTAATC-3’

Human IFNg Forward: 5′-GATTTAGAGTAATTTGAA ATTTGTGG-3′

Human IFNg Reverse: 5′-CCTCCTCTAACTACT AATATTTATACC-3′

The PCR amplicon was cloned into a pGEMT easy vector (Promega) and then transformed into XL10-Gold ultracompetent bacteria (Agilent Technologies). Bacterial colonies were selected using a blue/white X-gal selection system after overnight growth, the cloning vector was then purified from individual colonies, and the genomic insert was sequenced. After bisulfite treatment, the methylated CpGs were detected as cytosines in the sequence, and unmethylated CpGs were detected as thymines in the sequence by using QUMA software (37).

WGBS was performed as described previously. Briefly, bisulfite-modified DNA sequencing libraries were generated using the EpiGenome kit (Epicentre) according to the manufacturer’s instructions. Bisulfite-modified DNA libraries were sequenced using Illumina HiSeq 4000 and NovaSeq 6000 systems (20). Sequencing data were aligned to the HG19 genome using the BSMAP v. 2.74 software (38). Differential analysis of CpG methylation among the datasets was determined with a Bayesian hierarchical model to detect regional methylation differences with at least three CpG sites (39).

### MiSeq

Naïve, endogenous LCMV-specific, and WT P14 LCMV-specific CD8 T cells were FACs purified from either WT B6 or Tet2cKO mice. DNA was isolated from the cells, bisulfite converted, and then PCR amplified using primers from the Integrated DNA Technologies custom oligos tool. Illumina overhang adapter sequences were added to each respective primer to make the products compatible with Illumina index and sequencing adapters. Amplified samples were analyzed using the MiSeq platform.

#### Mouse IFNg Forward

5’<u>TCGTCGGCAGCGTCAGATGTGTATAAGAGACAG</u>TATTTTTATTGTTGTGGTTGGTAGCT G-3’

#### Mouse IFNg Reverse

5’<u>GTCTCGTGGGCTCGGAGATGTGTATAAGAGACAG</u>CTTTCTTCTCCAAATTACTTTTAATC3’

### Generating human TET2-KO.B7-H3 CAR T cells

#### Generating B7-H3-CAR lentiviral vectors

Human B7-H3 CAR T cells were generated using the CAR backbone described in Nguyen et al 2020 (under revision). The vector was modified by removing the insulators from the self-inactivating (SIN) 3’ partially-deleted viral LTRs (40,41). Briefly, codon-optimized DNA encoding B7-H3 specific scFv derived from the m276 monoclonal antibody (42) was synthesized by GeneArt (Thermo Fisher, Waltham, MA). The scFv was then ligated using infusion cloning (Takara Bio, Kusatsu, Shiga, Japan) into the backbone encoding a CD8α transmembrane domain, CD28 costimulatory domain, and CD3ζ activation domain. All final constructs were verified by sequencing. High titer lentiviral particles were then generated by the St. Jude Vector Core as described in (43).

#### TET2 Knock-out T cells

*TET2* and control knock-out T cells were generated using CRISPR-Cas9 technology. At 48 hours post-activation, sorted naïve T cells were nucleofected with *TET2* or control (mCherry) sgRNAs as Cas9 ribonucleoprotein (RNP) complexes. We used a *TET2* sgRNA (5’-CGAAGCAAGCCTGATGGAACNGG-3’, Synthego, Menlo Park, CA) and a control sdRNA targeting *mCherry* (5’ – CAAGUAGUCGGGGAUGTCGG -3’, Synthego, Menlo Park, CA). RNPs were pre-complexed at a sgRNA:Cas9 ratio of 4.5:1, prepared by adding 3 μL of 60 μM sgRNA (Synthego, Menio Park, CA) to 1 μL of 40 μM Cas 9 (Macro Lab, University of California, Berkeley), incubated at room temperature for 10 min and then stored at -20 ° C for later use. Activated T cells were collected by gentle pipetting up and down and then pelleted at 0.5×10^6^ cells per reaction. Cell pellets were resuspended in 17 μL of P3 transfection solution (13.94 μL of Nucleofector Solution with 3.06 μL of supplement, Lonza, Walkersville, MD) and 4 μL of RNP complexes were added for each reaction. Electroporation was then performed by transferring 20 μL of T cell-RNP mixture into the transfection vessel and using the Lonza transfection program EH:115 (4D-Nucleofector, Lonza, Walkersville, MD). Electroporated T cells were then allowed to recover for overnight in RPMI media supplemented with 20% FBS and 1% glutamax and in the presence of IL-7 and IL-15 cytokines at 10ng/mL and 5ng/mL respectively.

#### Transducing CAR T cells

For generating human CAR T cells, electroporated cells were collected after 24 hours and washed with complete RPMI media (10% FBS with 1% Glutamax). Cells were then plated at 0.5×10^6^cells/well in 500 μL of complete RPMI media supplemented with IL-7 and IL-15 cytokines. T Cells were transduced by adding the lentiviral particles at a multiplicity of infection of 50, and protamine sulfate at 8ug/ml. Transduced T cells were then expanded until day 10 post-transduction with frequent supplementation with fresh media containing IL-7 and IL-15. CAR detection was performed using anti-Fab specific antibody at day 5 post-transduction (109-606-006, Jackson ImmunoResearch, West Grove, PA).

### TET2-KO.B7-H3 CAR T cell activation assay

To evaluate the interaction of signal 3 cytokine signaling and DNA methylation in TET2-KO CAR T cells, transduced cells were activated with plate-bound recombinant human B7-H3 protein. Briefly, non-tissue culture plates were coated overnight at 4゜C with recombinant human B7-H3 protein (R&D Systems, Minneapolis, MN) at 10 μg/well in 500 μL of PBS. Wells coated with PBS only served as unstimulated controls. TET2 and control knockout CAR T cells were then washed with complete RPMI media and added to coated wells at 1×10^6^ cells/well in the presence or absence of IL-12 (10ng/mL). Stimulated and control unstimulated CAR T cells were then collected at day 14 post-activation for analysis.

## ACKNOWLEDGMENTS

We would like to thank Drs. Yiping Fan and Jeremy Crawford for bioinformatic analysis of methylation profiling. This work was supported by the National Institutes of Health (R01AI114442 and R01CA237311 to BY, loan repayment program to CZ) and the American Lebanese Syrian Associated Charities (ALSAC to BY).

## Figure Legends

**Supplemental figure 1.**
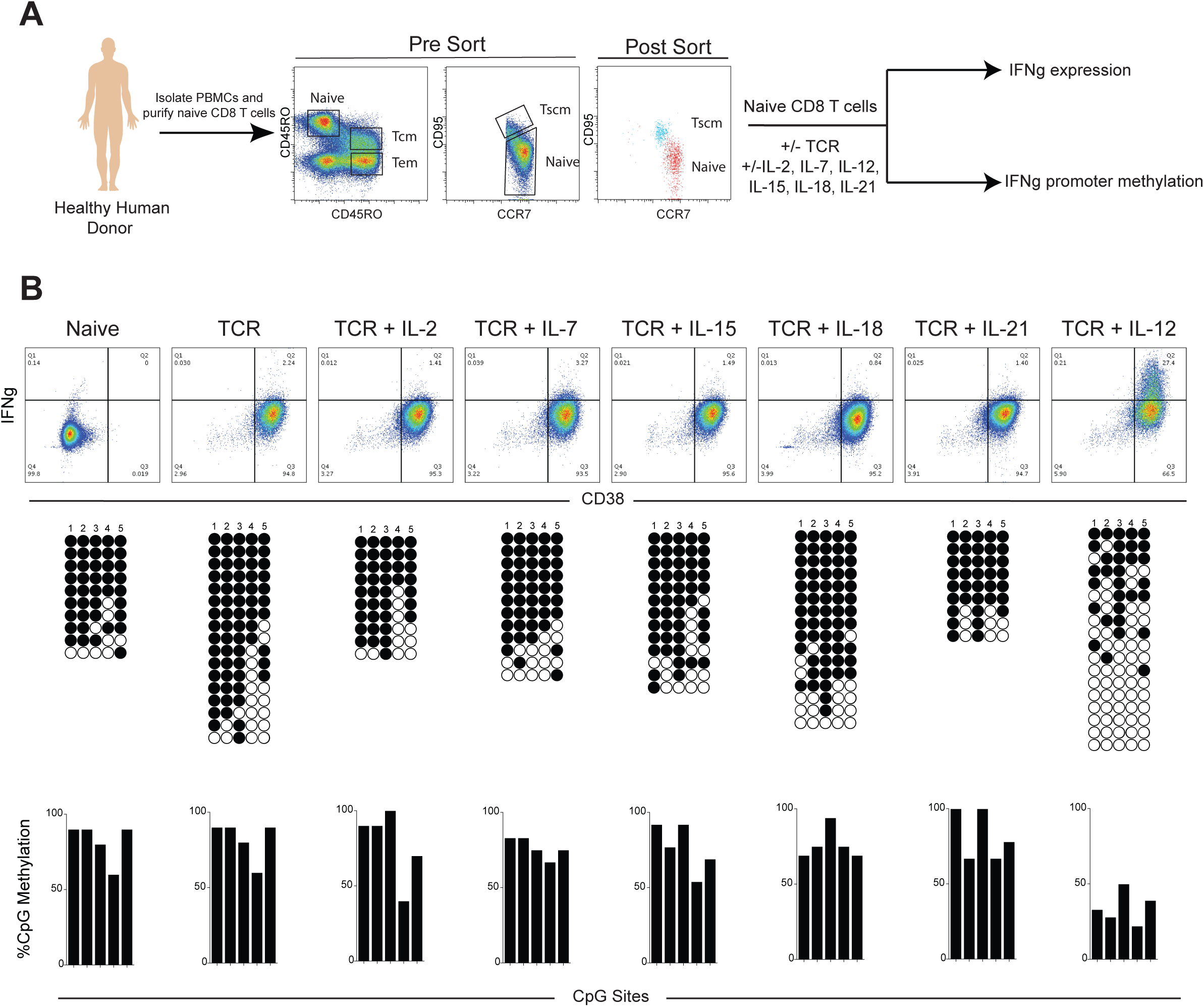
*Ex vivo* screening for signal 3 cytokines that promote IFNg expression and locus demethylation. **(A)** Experimental setup showing isolation of naïve CD8 T cells for *in vitro* stimulation with or without anti CD3/CD28 (TCR) and with or without the individual cytokines IL-2, IL-7, IL-12, IL-15, IL-18, IL-21 added once on day 0. **(B)** CD38 and IFNg expression of naïve, TCR stimulated, and TCR + cytokine after 7 days of *ex vivo* culture. *Lower panel:* Representative IFNg promotor bisulfite sequencing methylation profiles and summary graphs.

**Supplemental figure 2.**
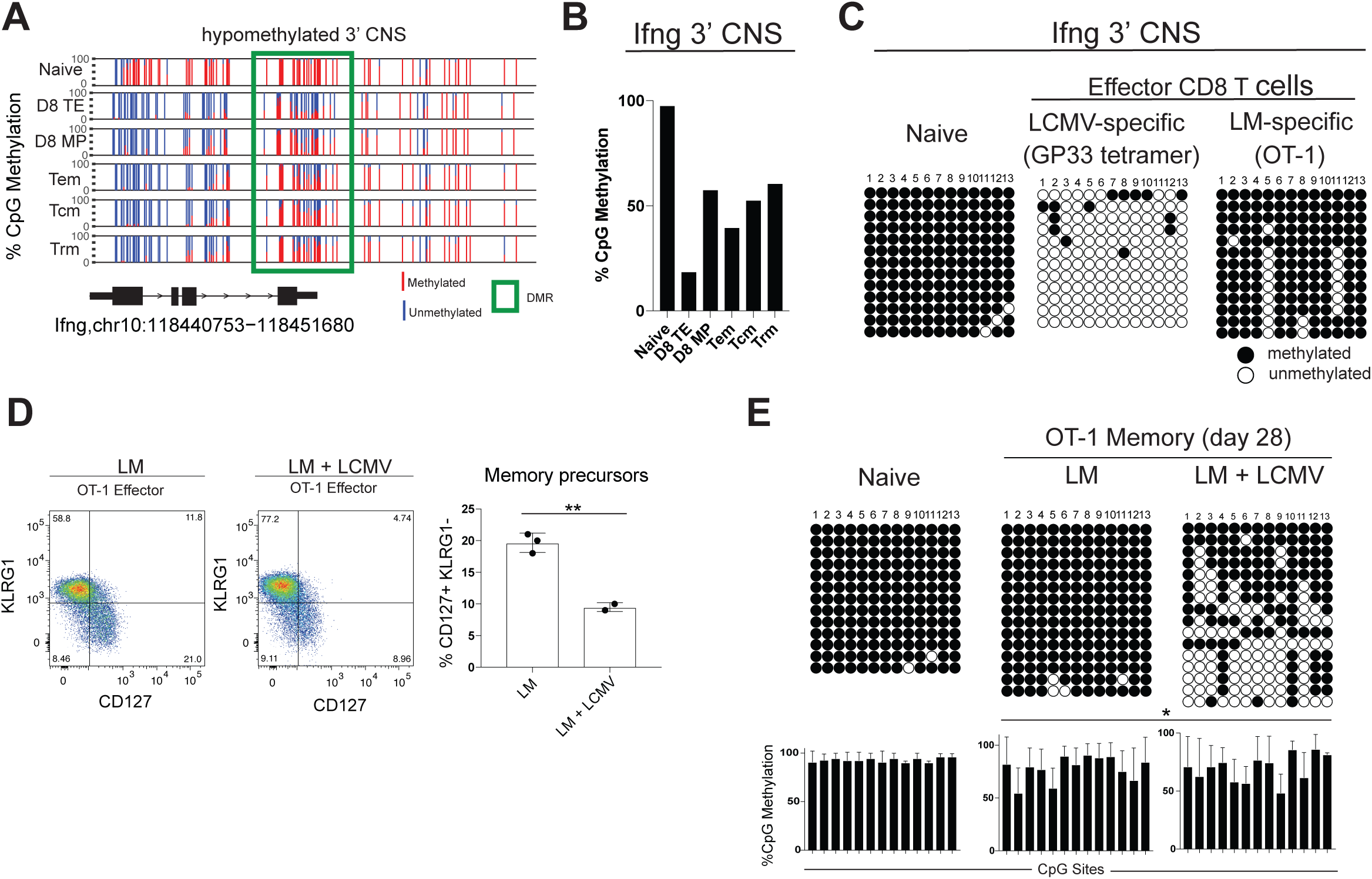
Infection with LCMV Armstrong results in Ifng intergenic demethylation variability in LCMV-specific effector CD8 T cells. **(A)** Methylation profiles of the IFNg gene from naïve, day 8 terminal effector (TE), day 8 memory precursor (MP), effector memory (Tem), central memory (Tcm), and tissue resident memory (Trm) P14 CD8 T cell published whole genome DNA methylation datasets. Memory profiles were generated from P14 cells after 30 days post LCMV Armstrong infection. Highlighted DMR overlaps with vertebrate conserved non-coding sequence (CNS). **(B)** Summary of total methylation identified in highlighted DMR from panel A. **(C)** Representative targeted bisulfite sequencing methylation profiles of naïve CD8 T cells and LCMV-specific and LM-specific effector CD8 T cells obtained 8 days post acute infection. For all representative bisulfite sequencing analysis, each horizontal line represents a clone and each vertical line represents a different CpG site. Black circles represent methylated CpGs while white circles represent unmethylated CpGs. **(D)** Representative phenotypic analysis and summary graph of OT1 cells isolated from the spleen of LM infected or LM + LCMV coinfected mice at day 8 post infection. **(E)** Representative and summary graphs of IFNg CNS bisulfite sequencing methylation profiles of naïve CD8 T cells and LM-specific and LCMV-specific effector CD8 T cells at day 8 post infection. *Lower panel*: Bar graphs showing % CpG methylation (mean ± SEM) at each individual CpG site of the IFNg locus. n=3 for LM and n=2 for LM+LCMV summary graphs.

